# Identification of *kit-ligand a* as the Gene Responsible for the Medaka Pigment Cell Mutant *few melanophore*

**DOI:** 10.1101/713107

**Authors:** Yuji Otsuki, Yuki Okuda, Kiyoshi Naruse, Hideyuki Saya

## Abstract

The body coloration of animals is due to pigment cells derived from neural crest cells, which are multipotent and differentiate into diverse cell types. Medaka (*Oryzias latipes*) possesses four distinct types of pigment cells known as melanophores, xanthophores, iridophores, and leucophores. The *few melanophore* (*fm*) mutant of medaka is characterized by reduced numbers of melanophores and leucophores. We here identify *kit-ligand a* (*kitlga*) as the gene whose mutation gives rise to the *fm* phenotype. This identification was confirmed by generation of *kitlga* knockout medaka and the findings that these fish also manifest reduced numbers of melanophores and leucophores and fail to rescue the *fm* mutant phenotype. We also found that expression of *sox5*, *pax7a*, *pax3a*, and *mitfa* genes is down-regulated in both *fm* and *kitlga* knockout medaka, implicating c-Kit signaling in regulation of the expression of these genes as well as the encoded transcription factors in pigment cell specification. Our results may provide insight into the pathogenesis of c-Kit–related pigmentation disorders such as piebaldism in humans, and our *kitlga* knockout medaka may prove useful as a tool for drug screening.

## INTRODUCTION

The body coloration of animals is attributable to pigment cells in the skin that are derived from neural crest cells and which provide protection from ultraviolet light as well as play a role in sexual selection and mimesis. Whereas mammals and birds possess a single type of pigment cell known as a melanocyte, six types of pigment cells known as chromatophores—melanophores (black), xanthophores (yellow), iridophores (iridescent), erythrophores (red), cyanophores (blue), and leucophores (white)—have been identified in fish (Fujii 1993). Given that these pigment cells are all derived from neural crest cells and can be readily distinguished on the basis of their color, fish have been studied as model organisms for characterization of the mechanisms underlying regulation of cell fate determination in multipotent cells. Medaka (*Oryzias latipes*) possesses four of these chromatophore types: melanophores, xanthophores, iridophores, and leucophores (Takeuchi 1976; Kelsh *et al*. 1996; Kelsh *et al*. 2004).

Intermediate progenitors and key transcription factors required for fate specification in neural crest cells have been identified (Bhatt *et al*. 2013). Although the molecular mechanisms of melanophore differentiation in fish have been relatively well characterized, those underpinning the differentiation of other pigment cell types have remained largely unknown. Characterization of the molecular mechanisms responsible for abnormal body coloration is expected to provide insight into the development of chromatophores, with such mutants also being applicable to the screening of drugs and studies of regenerative medicine related to skin pigmentation disorders in humans.

Various medaka mutants with abnormal body coloration have been described (Tomita 1992), and causal genes for such mutants have been identified. Such medaka mutant collections provide an important resource for studies of the genetic basis of fate determination in neural crest cells. One such recessive mutant, *few melanophore* (*fm*), is characterized by reduced numbers of melanophores and leucophores (Kelsh *et al*. 2004). The causal gene for this mutant has remained unknown, but its identification would be expected to provide insight into the differentiation of these two pigment cell types.

The *pax7a* gene has been implicated in fate specification of a shared, partially restricted progenitor of the xanthophore and leucophore lineages in medaka (Kimura *et al*. 2014), and *sox5* functions in a cell-autonomous manner to control the specification of xanthophores from the shared xanthophore-leucophore progenitor (Nagao *et al*. 2014). We have here adopted genetic approaches to identify the causal gene and molecular mechanisms underlying the phenotype of *fm* medaka. We found that the *fm* locus includes a mutated version of the *kit-ligand a* gene (*kitlga*, DK099743) and that expression of *pax7a*, *sox5*, *pax3a*, and *mitfa* is down-regulated in both *fm* and *kitlga* knockout (KO) medaka. Our results thus suggest that the abnormal coloration of *fm* medaka is caused by disruption of the *kitlga* gene.

## MATERIALS AND METHODS

### Medaka strains and maintenance

The *fm* strain (ID: MT48) of medaka has been described previously (Kelsh *et al*. 2004). The Sakyo strain (ID: WS1164) is normal with regard to the production of all four pigment cell types and was thus studied as the wild type (WT). The Kaga strain (ID: IB833) was used for crossing in genetic mapping. All medaka strains were obtained from NBRP Medaka (https://shigen.nig.ac.jp/medaka/top/top.jsp). Fish were maintained in a recirculating system with a 14-h-light, 10-h-dark cycle at 28°C. All medaka experiments were performed according to a protocol approved by the Animal Care and Use Committee of Keio University (permit no. 16041).

### Observation of pigment cells

Larvae and adult fish were anesthetized with tricaine mesylate, and dorsal body images were obtained at a constant magnification and resolution (1280 × 968 pixels) with a Leica MZ12.5 stereomicroscope or a Nikon SMZ25 microscope. For counting the number of melanophores, scales of adults were treated with epinephrine (2 mg/ml) to induce melanin aggregation. Melanophores and leucophores on the dorsal side of each larva were counted at 3 days posthatching (dph), and embryos were also evaluated at stages 26, 30 and 36. The total area of leucophores of each embryo at stage 30 and the area of individual melanophores at stage 36 were measured with Image J software (Schneider *et al*. 2012). Xanthophores were counted in larvae and on the scales of adult fish observed under ultraviolet light after treatment with 10uM melatonin for 10min and fixation with 10% paraformaldehyde. Iridophores were evaluated on the basis of iris luminosity and with the use of Image J software (Schneider *et al*. 2012).

### Positional cloning of the gene mutated in *fm* medaka

Crossing of the F_1_ generation obtained by breeding the *fm* mutant with the Kaga strain yielded 156 F_2_ offspring with the *fm* phenotype and 30 siblings with the WT phenotype, which were collected and subjected to bulk segregation analysis with the M-marker 2009 system as described previously (Kimura and Naruse, 2010). For further recombination analysis, polymorphic markers were isolated with reference to the medaka genome database (http://medakagb.lab.nig.ac.jp/Oryzias_latipes/index.html). Detailed information on the markers, including primers for polymerase chain reaction (PCR) amplification and restriction enzymes for genotyping, is provided in Table 1.

**Table 1.**
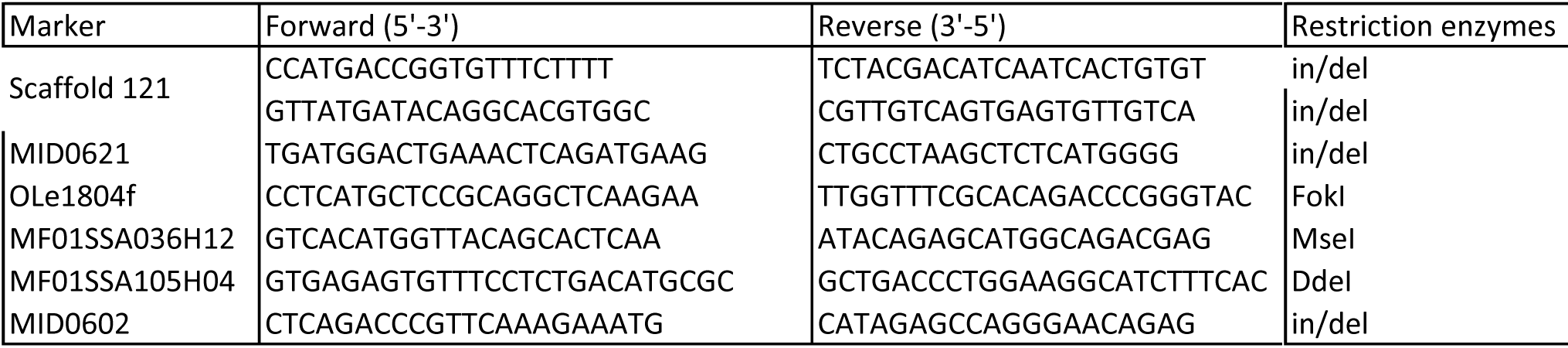
Primer sequences and restriction enzymes for genotyping

### 5’-RACE analysis

Body tissue of medaka at 3 dph was minced and then processed with a RNeasy Minikit (Qiagen) for extraction of total RNA. The RNA was subjected to reverse transcription (RT) for 15 min at 37°C with the use of a PrimeScript RT Reagent Kit with gDNA Eraser (Takara), after which the reaction was terminated by incubation at 85°C for 5 s. The resulting cDNA was subjected to 5’ rapid amplification of cDNA ends (5’-RACE) with the use of a GeneRace Kit (Invitrogen) and with region-specific primers.

### Generation of *kitlga* KO medaka

Gene targeting with the CRISPR/Cas9 system was performed as described previously (Ansai and Kinoshita 2014). The single guide RNAs (sgRNAs) were designed to target exons 2 and 4 of *kitlga* (see Figure 5A), and microinjection was performed with the Sakyo strain. Genomic DNA was purified from fin clips. For sequencing, exons 2 to 4 of *kitlga* were amplified with the use of the Ampdirect reagent and BIOTAQ HS DNA polymerase (Shimadzu). The PCR incubation protocol included an initial incubation at 95°C for 10 min; 35 cycles at 95°C for 30 s, 68°C for 30 s, and 72°C for 90 s; and a final incubation at 72°C for 10 min. The PCR products were separated by electrophoresis, purified with the use of a QIAquick Gel Extraction Kit (Qiagen), and sequenced. The PCR primers for amplification and sequencing were KLG ex2-F (5’-TGATCTTAGTCATGTTTTT-3’;) and Crispr-R (5’-AGCAGCACATGGACTTATTCC-3’).

### Genotyping of *fm* medaka

Genomic DNA was extracted from fin clips that were prepared from anesthetized fish and fixed in 100% methanol. The samples were suspended in 100 µl of lysis buffer [20 mM Tris-HCl (pH 8.0), 5 mM EDTA, 400 mM NaCl, 0.3% SDS, and proteinase K (10 mg/ml)], incubated at 56°C for at least 2 h, and then stored at –80°C until analysis. They were subsequently applied directly to a PCR reaction mixture containing Ampdirect (Shimadzu). The PCR conditions for *fm* medaka included an initial incubation at 95°C for 10 min; 35 cycles at 95°C for 30 s, 56°C for 30 s, and 72°C for 60 s; and a final incubation at 72°C for 10 min. The primers were KLG ex2-F (5’-TGATCTTAGTCATGTTTTT-3’), fm ex2-R (5’-TGTGTCACTAACTACAGCATCT-3’), KLG ex5-F (5’-GTTATGATACAGGCACGTGGC-3’), and KLG ex5-R (5’-ACTGTTGTGAGTGACTGTTGC-3’).

### RT and real-time PCR analysis

Total RNA was extracted and subjected to RT as described for 5’-RACE. The resulting cDNA was subjected to real-time PCR analysis with the use of a Thermal Cycler Dice Real Time System (TAKARA BIO INC.). The PCR primers for medaka *kitlga*, *pax7a*, *sox5*, *pax3a*, and *mitfa* are listed in Table 2. The abundance of each target mRNA was normalized by that of elongation factor (EF)–1α mRNA as an invariant control.

**Table 2.**
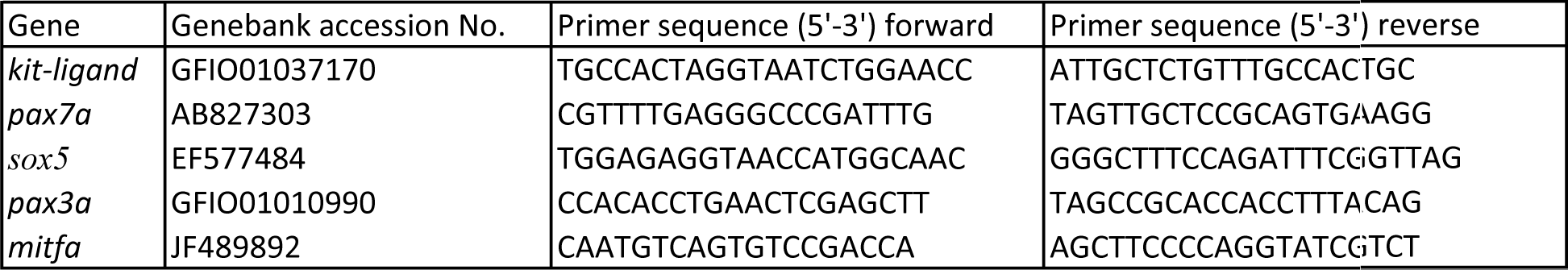
Primer sequences for quantitative RT-PCR.

### Phylogenetic analysis of *kitlg* genes in teleosts

A phylogenetic tree for *kitlg* genes was generated by the maximum likelihood method on the basis of amino acid sequences listed in Table S1 and with the use of MEGA-X software (Kumar *et al*. 2018).

### Statistical analysis

Quantitative data are presented as means + 95% confidence interval and were compared among groups by one-way analysis of variance (ANOVA) followed by Tukey’s post hoc test. A *p* value of <0.05 was considered statistically significant.

### Data availability

All medaka strains are available from NBRP Medaka (https://shigen.nig.ac.jp/medaka/top/top.jsp). Supplemental files available at FigShare. Sequence data are available at GenBank; the accession numbers are listed in Table S1 and 2.

## RESULTS

### The *fm* medaka mutant has reduced numbers of melanophores and leucophores

*fm* medaka is a spontaneous mutant discovered by Takahashi in 1965 and established by Tomita in 1971 (Tomita 1975). It was first described as having a reduced number of melanophores, but was later shown by Kelsh et al. (2004) to also be characterized by a reduced number of leucophores and an abnormal shape of melanophores. We first examined the numbers of melanophores, leucophores, and iridophores (determined by iris luminosity measurement) in embryos, larvae, and adults of WT, *fm* heterozygous (*fm het*), and *fm* medaka. Xanthophores were also examined in the scales of adult fish as well as on the lateral side of larvae examined under ultraviolet light.

Melanophores were apparent on the dorsal side and yolk sphere and leucophores were detected on the dorsal side of the head in WT embryos at stage 26 (Figure 1A). Iridophores could not be examined because eyes were not silver at this stage. In contrast to WT and *fm het* medaka at this stage, melanophores were not found in the center of the head in *fm* embryos (Figure 1A–C). At stage 30, melanophores had spread throughout the lateral side of the back and their number had increased in WT and *fm het* medaka, whereas their number remained low in *fm* embryos (Figure 1D–F). The area of leucophore pigmentation was also narrower in *fm* medaka than in WT and *fm het* embryos (Figure 1D–F). At stage 36, many melanophores were visible throughout the entire body, with leucophores coexisting with melanophores along the back, of WT and *fm het* embryos (Figure 1G, H). The numbers of melanophores and leucophores in *fm* had remain lower than those in WT or *fm het* medaka (Figure 1G–I). Quantitative analysis revealed that the numbers of both melanophores and leucophores were significantly lower in *fm* embryos than in WT or *fm het* embryos from stages 26 to 36 (Figure 1J, K). It was difficult to count the number of leucophores at stage 30, but the total area of these cells was significantly lower in *fm* embryos than in WT or *fm het* embryos at this stage (Figure 1K). Moreover, melanophores appeared smaller in *fm* medaka than in WT and *fm het* embryos at stage 36 (Figure 1L). The luminosity value of iridophores at stage 30 or 36 did not differ among the three genotypes (Figure 1M).

**Figure 1.**
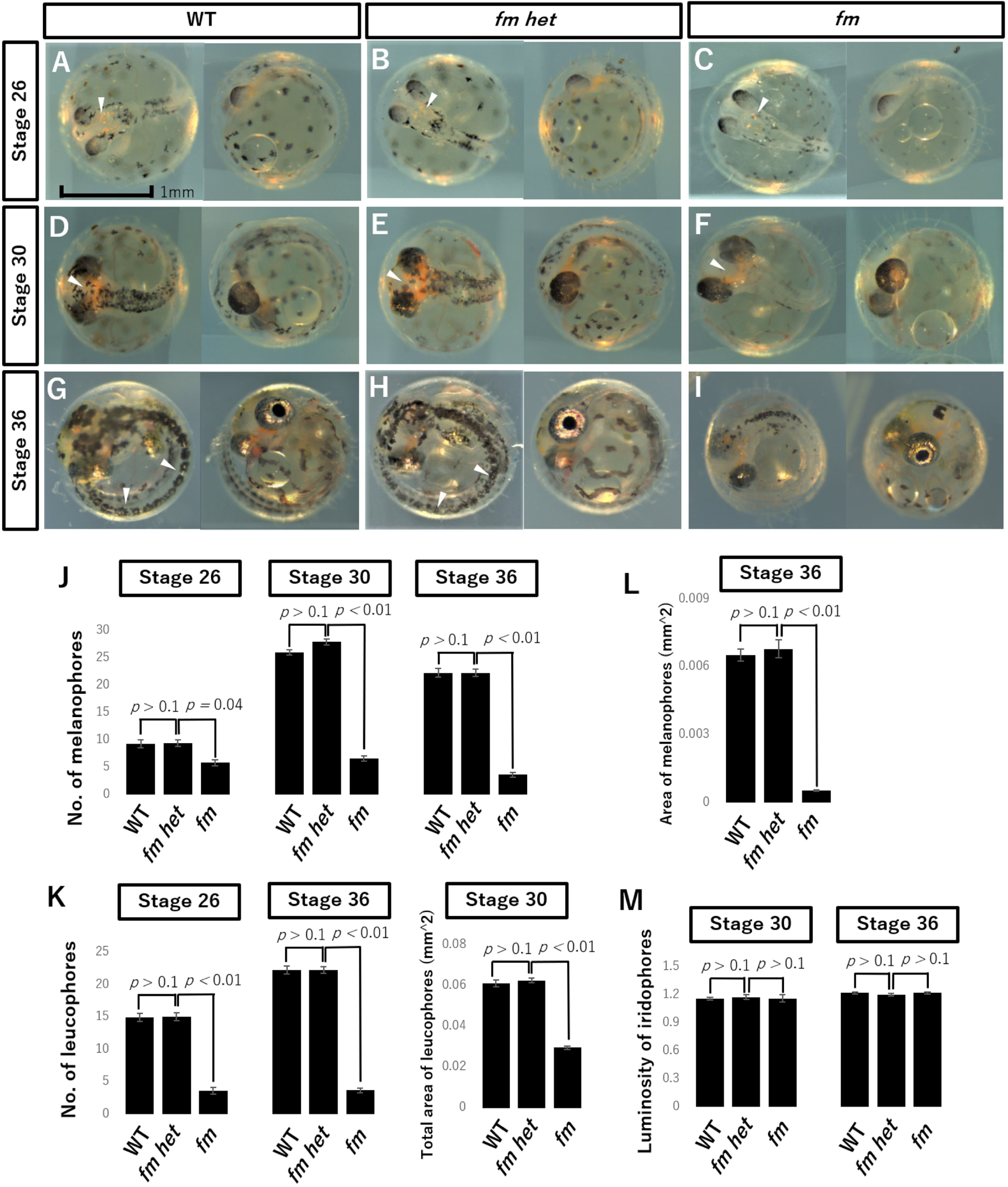
The numbers of melanophores and leucophores are reduced in embryos of *fm* medaka. (A–I) Photographs of WT (A, D, G), *fm het* (B, E, H), and *fm* (C, F, I) medaka embryos at stage 26 (A–C), stage 30 (D–F), or stage 36 (G–I). Arrowheads indicate a reduced number of leucophores on the dorsal head (A–C), a narrower area of leucophore pigmentation (D–F), or stage 36 (G–I) in *fm* medaka compared with WT or *fm het* embryos. Scale bar, 1 mm. (J) Number of melanophores in embryos of the three genotypes at stages 26, 30, and 36. (K) Number or total area of leucophores at stages 26 and 36 or at stage 30, respectively. (L) Area of individual melanophores at stage 36. (M) Luminosity value of iridophores at stages 30 and 36. All quantitative data are means + 95% confidence interval (*n* =15 larvae for each genotype). The *p* values were determined by one-way ANOVA followed by Tukey’s post hoc test.

Examination of WT and *fm het* larvae at 3 dph revealed that most melanophores colocalized with leucophores on the dorsal side and in the head region (Figure 2A). In *fm* medaka, although the differentiation of all chromatophores was apparent, and melanophores and leucophores were also positioned at the dorsal midline, the melanophores appeared smaller than in WT and *fm het* larvae (Figure 2A). Furthermore, whereas the numbers of melanophores and leucophores had increased to ~25 in the dorsal midline of the trunk in WT and *fm het* larvae at 3 dph, those in the *fm* mutant remained significantly smaller (Figure 2B). There was no apparent difference in the number of xanthophores on the lateral side of larvae examined under ultraviolet light (Figure 2C). The luminosity value of iridophores in the iris of larvae at 3 dph also did not differ significantly among the three genotypes (Figure 2D, E).

**Figure 2.**
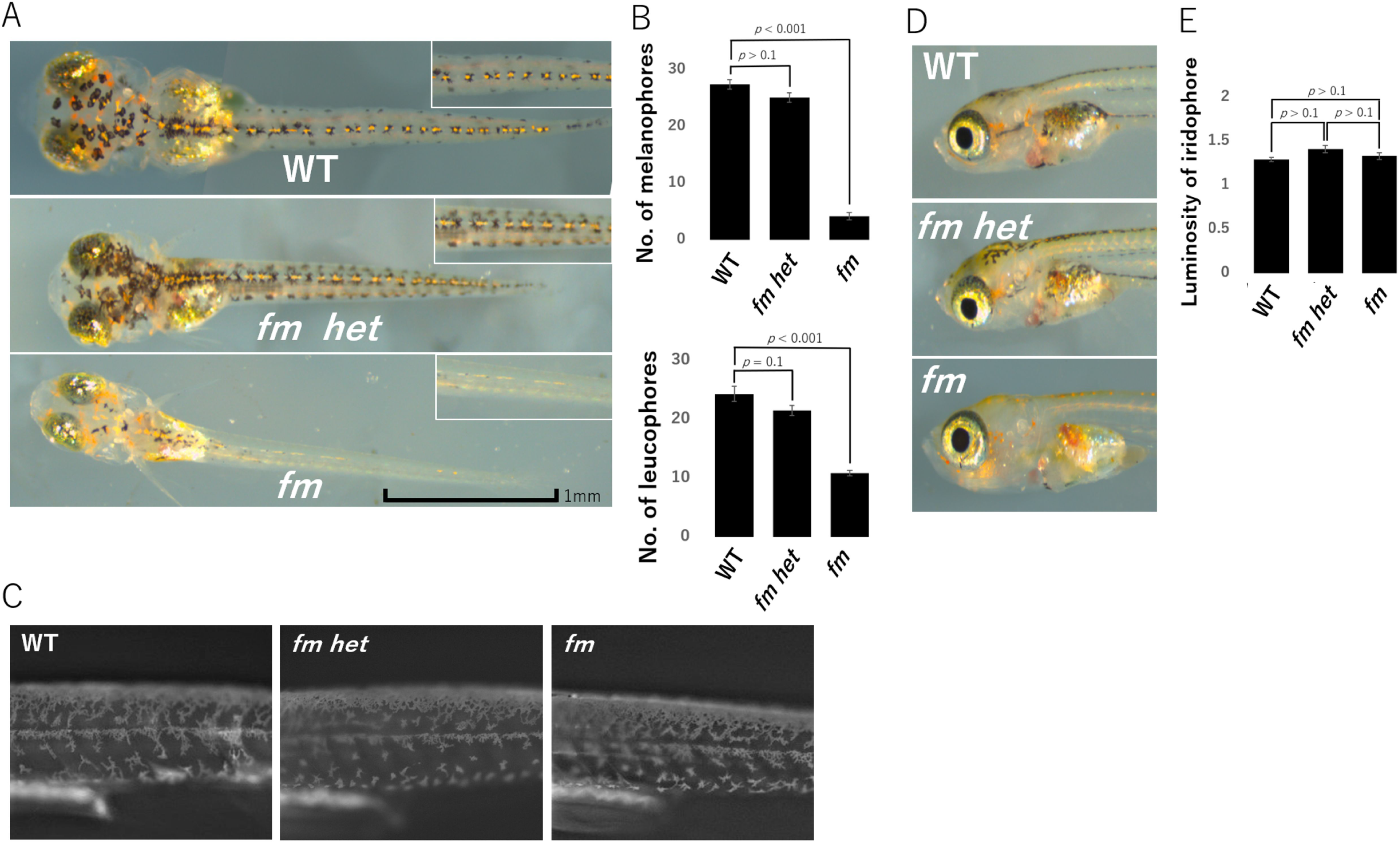
Reduced numbers of melanophores and leucophores in *fm* medaka larvae at 3 dph. (A) Photographs of the dorsal side of WT, *fm het*, and *fm* larvae. Insets show higher magnification views. Scale bar, 1 mm. (B) Numbers of melanophores and leucophores on the dorsal side of the body for larvae of the three genotypes. (C) Photographs of the lateral side of larvae under ultraviolet illumination. (D) Photographs of the lateral side of larvae showing the iris. (E) Luminosity value of iridophores in the iris. All quantitative data are means + 95% confidence interval (*n* =15 larvae for each genotype). The *p* values were determined by one-way ANOVA followed by Tukey’s post hoc test.

The back of adult WT and *fm het* medaka appeared black as a result of the large number of melanophores, whereas the *fm* mutant appeared paler because of the continued reduction in melanophore number (Figure 3A). The numbers of melanophores and leucophores on scales were also larger for WT and *fm het* adults than for *fm* adults (Figure 3B, C). The number of xanthophores on scales did not differ significantly among adults of the three genotypes (Figure 3B, D). The luminosity value of iridophores in the iris was also similar for adults of all three genotypes (Figure 3E, F). As with embryos and larvae, there were no apparent differences in chromatophores between WT and *fm het* adults, consistent with the notion that *fm* is a recessive mutation that reduces the numbers of melanophores and leucophores specifically.

**Figure 3.**
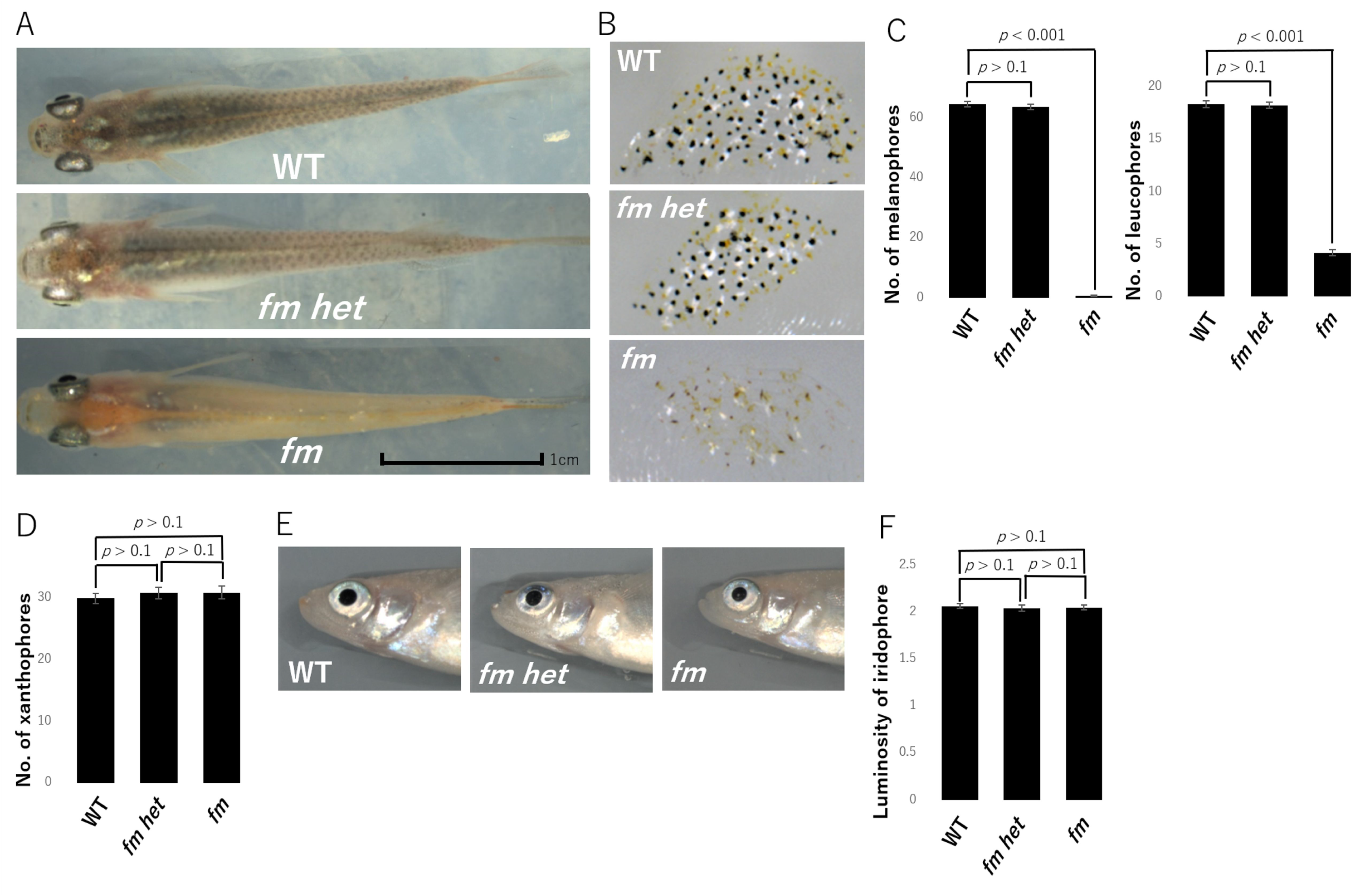
Reduced numbers of melanophores and leucophores in adult *fm* medaka. (A) Photographs of the dorsal side of adult WT, *fm het*, and *fm* medaka. Scale bar, 1 cm. (B) Photographs of the scales of adult medaka of the three genotypes. (C) Numbers of melanophores and leucophores for scales on the dorsal side of adult medaka. (D) Number of xanthophores for scales on the dorsal side of adult medaka. (E) Photographs of the lateral side of adult medaka showing the iris. (F) Luminosity value of iridophores in the iris of adult medaka. All quantitative data are means + 95% confidence interval (*n* = 15 adults for each genotype). The *p* values were determined by one-way ANOVA followed by Tukey’s post hoc test.

### The *fm* locus contains the *kitlga* gene

To identify the *fm* locus, we adopted a positional cloning approach. Bulk segregation analysis with M-marker 2009 (Kimura and Naruse 2010) suggested that the *fm* locus was present in linkage group 6. Analysis of linkage among the *fm* locus and DNA markers in linkage group 6—including MID0602, MID0621, OLe1804f, MF01SSA036H12, and MF01SSA105H04 (Naruse *et al*. 2004; Kimura and Naruse 2010)—confirmed that *fm* maps to chromosome 6 (Figure 4A).

**Figure 4.**
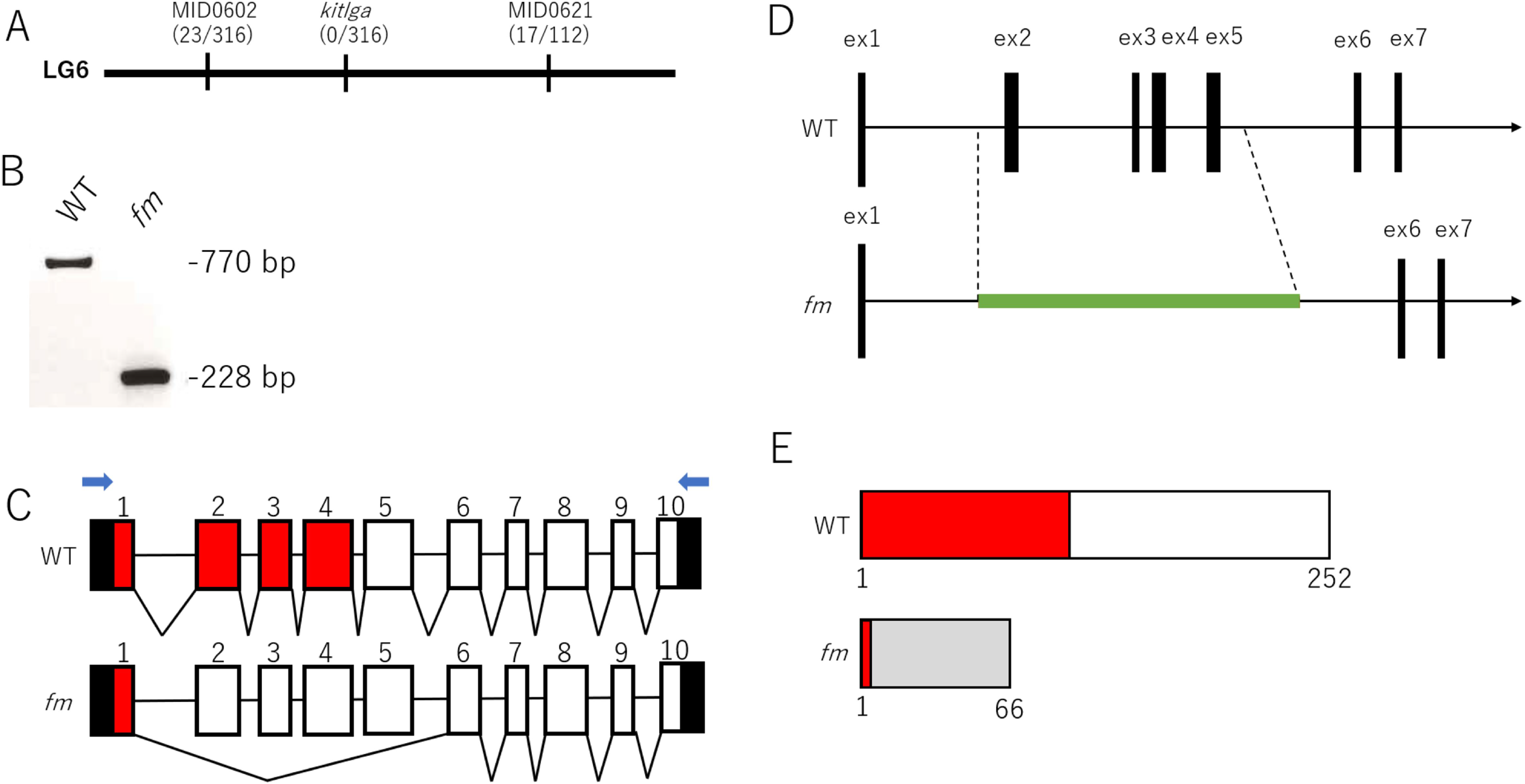
Mapping of the *fm* locus. (A) The *fm* locus was mapped to the region between MID0602 and MID0621 in linkage group 6 (LG6). A *kitlga* gene marker did not show any recombination with the *fm* phenotype (indicated by the numbers in parentheses). (B) RT-PCR analysis of total RNA from WT and *fm* medaka with the primer set indicated by the blue arrows in (C). (C) The *kitlga* gene of WT medaka comprises 10 exons (numbered boxes). Sequencing of cDNA generated by 5’-RACE showed that exons 2 to 5 are skipped in the *fm* mutant. The 5’ and 3’ untranslated regions are shown in black, and exonic sequence encoding the SCF domain in red. (D) Sequencing of genomic DNA revealed that exons 2 to 5 (3.6 kb) of *kitlga* are replaced with a 3.9-kb sequence (green line) in the *fm* genome. (E) The WT *kitlga* gene encodes a 252–amino acid protein containing an SCF domain (red box). The *fm* mutation is predicted to result in the generation of a truncated protein, with the gray box representing the altered frame.

We next focused on the c-Kit signaling pathway given that c-Kit receptor mutants of zebrafish and guppy (Kelsh *et al*. 2004; Kottler *et al*. 2013) show marked similarity to the *fm* medaka mutant. In particular, the embryonic phenotype of the *fm* mutant, characterized by a reduced number and smaller size of melanophores, was found to be highly reminiscent of that of the zebrafish *sparse/kit* mutant (Kelsh *et al*. 2004). We searched the genomic regions of *kita* (*kit receptor a*) and *kitb* (*kit receptor b*) in the Ensembl database? and found that *kita* is located on chromosome 4 and *kitb* on chromosome 1 of the medaka genome. An Ensembl-based search for the map position of the gene encoding Kit ligand (*kitlg*) revealed that this gene is located in scaffold121, which had not been mapped to a chromosome in MEDAKA1 (Ensembl release 93). The *kitlg* gene was subsequently mapped to the region spanning 2469132 to 2512640 bp of chromosome 6 (Ensembl genome assembly ASM223467v1). A second *kitlg* gene was also identified on chromosome 23, however. To examine the relation between these two *kitlg* genes, we constructed a phylogenetic tree of teleost *kitlg* genes based on amino acid sequences shown in Table S1. The *kitlg* gene on chromosome 6 of medaka was thus found to belong to the *kitlga* clade and that on chromosome 23 to the *kitlgb* clade (Figure S1). We therefore designated these two medaka *kitlg* genes as *kitlga* and *kitlgb*, respectively. We designed a *kitlga* gene marker and found that the gene maps to chromosome 6 between MID0602 and MID0621 and that there was no recombination between the *kitlga* gene marker and the *fm* phenotype (Figure 4A).

To determine whether the *fm* mutant harbors a mutation in *kitlga*, we performed RT-PCR analysis. Such analysis revealed deletion of a portion of *kitlga* cDNA in the mutant (Figure 4B). Analysis by 5’-RACE identified a 475-bp deletion corresponding to skipping of exons 2 to 5 (Figure 4C). Sequencing of this genomic region of *fm* medaka revealed substitution of the 3.6-kb region encompassing exons 2 to 5 of the WT gene with a 3.9-kb sequence of unknown origin (Figure 4D). The medaka *kitlga* gene comprises 10 exons with a 756-bp open reading frame that encodes a 252–amino acid protein. The *fm* mutation results in the generation of an open reading frame for a truncated protein that lacks the stem cell factor (SCF) domain and would therefore be expected to be nonfunctional (Figure 4E). A BLASTX analysis of the genomic sequence of the mutated *kitlga* gene in *fm* medaka revealed that the insertion shows marked sequence similarity to the transposase encoded by the transposon Caenorhabditis briggsae 1 (Tcb1), which has been identified in the genomes of other fish species such as zebrafish and rainbow trout. Moreover, we found that this transposon-like sequence is also present at >100 additional regions of the current Ensembl genome assembly (for medaka ASM223467v1). These results suggested that the phenotype of *fm* medaka is attributable to insertion of a Tcb1-like transposon at the *kitlga* locus.

### CRISPR/Cas9–mediated knockout of the *kitlga* gene induces an *fm*-like phenotype

To confirm *kitlga* as the causal gene of the *fm* mutant, we generated *kitlga* KO medaka with the use of the CRISPR/Cas9 system and sgRNAs targeted to the splice donor sites of exons 2 and 4 (Ansai and Kinoshita 2014). Microinjection of one cell–stage WT embryos resulted in the generation of some larvae with reduced numbers of melanophores and leucophores at 3 dph, a phenotype similar to that of the *fm* mutant. Control embryos injected with only sgRNA or Cas9 mRNA failed to give rise to larvae that mimicked the *fm* phenotype. We outcrossed the *kitlga* G_0_ medaka with WT fish to obtain F_1_ medaka, sequence analysis of which revealed that the CRISPR/Cas9 system induced a 9-bp deletion in exon 4 of *kitlga* that altered the amino acid sequence of the encoded protein (Figure 5A, B). We then generated homozygous *kitlga* KO medaka, which again manifested a phenotype indistinguishable from that of the *fm* mutant (Figure 5C, D). Given that the *kitlga* KO medaka were viable and fertile, we performed a complementation test to further verify that *kitlga* is the causal gene of the *fm* mutant. We obtained a total of 30 embryos from a cross between *kitlga* KO medaka and the *fm* mutant, with all larvae showing the same phenotype as the *fm* mutant (Figure 5C) characterized by reduced numbers of melanophores and leucophores (Figure 5D). Together, these results thus indicated that mutation of the *kitlga* gene is responsible for the *fm* phenotype of medaka.

### Expression of *pax3a*, *pax7a*, *sox5*, and *mitfa* is down-regulated in *fm* medaka

The *pax7a* gene is expressed in neural crest cells of medaka and functions as a molecular switch for the differentiation of multipotent progenitor cells into either xanthophores and leucophores or iridophores and melanophores, whereas the *sox5* gene is expressed in differentiating xanthophores and functions as a molecular switch in the specification of xanthophores versus leucophores (Kimura *et al*. 2014; Nagao *et al*. 2014). Sox5 belongs to the SOXD group of proteins and also plays a role in formation of the cephalic neural crest (Perez-Alcala *et al*. 2004). Pax3 and Pax7 are closely related transcription factors of the Pax family that manifest similar DNA binding activity in vitro, and Pax3 regulates the promoter of the Mitf (mouse microphthalmia-associated transcription factor) gene (Schäfer *et al*. 1994; Lacosta *et al*. 2005). To examine the molecular mechanisms underlying the reduction in the numbers of melanophores and leucophores in *fm* and *kitlga* KO medaka, we therefore determined the expression levels of *pax7a*, *sox5*, *pax3a*, and *mitfa*. RT and real-time PCR analysis revealed that the expression of each of these four genes was down-regulated in *fm* and *kitlga* KO larvae relative to WT larvae (Figure S2), suggesting that such down-regulation may contribute to the mutant phenotype.

**Figure 5.**
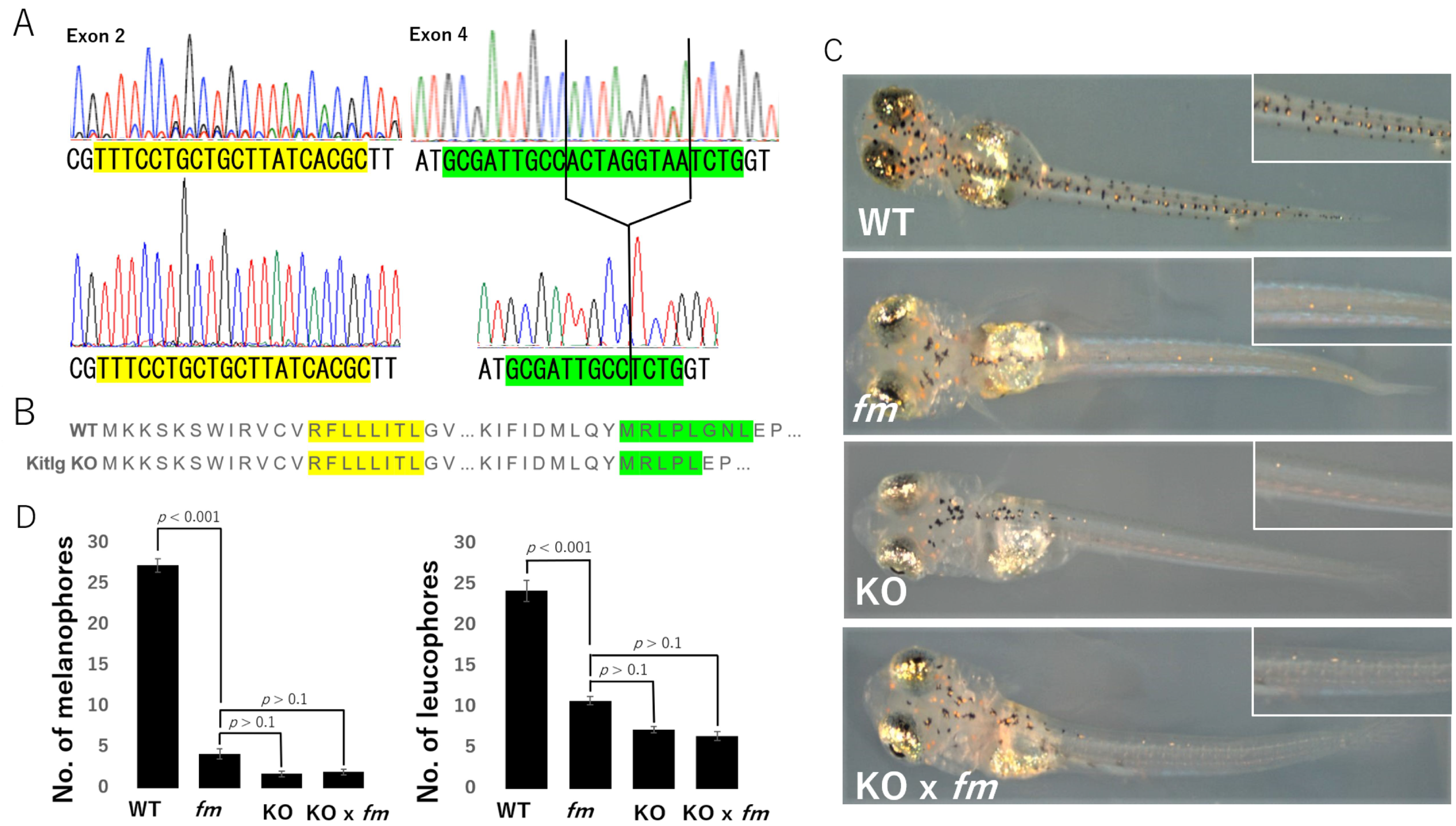
Targeting of the *kitlga* gene by the CRISPR/Cas9 system gives rise to an *fm*-like phenotype. (A) Sequencing of the CRISPR/Cas9 target region in manipulated medaka revealed a 9-bp deletion in exon 4 of *kitlga*. The upper and lower panels show the WT and mutated sequences of *kitlga*, respectively. Green, blue, black, and red traces indicate A, C, G, and T nucleotides, respectively. The sgRNA target sequences are shaded yellow and green. (B) Predicted amino acid sequences for *kitlga* in WT and *kitlga* KO medaka. The 9-bp deletion in exon 4 results in the deletion of three amino acids. (C) Photographs of WT, *fm*, and homozygous *kitlga* KO larvae at 3 dph as well as of a corresponding F_1_ larva derived from a cross between the *fm* mutant and *kitlga* KO medaka. Insets show higher magnification views. The body color of the *kitlga* KO medaka and of the cross between the *fm* and *kitlga* KO fish is similar to that of the *fm* mutant, indicating that the *kitlga* mutant could not rescue the *fm* phenotype. (D) Numbers of melanophores and leucophores on the dorsal side of the body of larvae as in (C). Data are means + 95% confidence interval (*n* = 15 larvae for each genotype). The *p* values were determined by one-way ANOVA followed by Tukey’s post hoc test.

## DISCUSSION

We have here identified *kitlga* as the gene responsible for the *fm* mutant of medaka, which is characterized by reduced numbers of melanophores and leucophores in embryos, larvae, and adult fish. Moreover, genomic PCR, RT-PCR, and 5’-RACE analyses revealed that the *fm* mutation is a deletion of exons 2 to 5 of *kitlga* and their replacement with a transposon-like sequence, which likely gives rise to a null allele of *kitlga*. Larvae of *kitlga* KO medaka established with the CRISPR/Cas9 system also manifested reduced numbers of melanophores and leucophores, and the progeny of a cross between *fm* and *kitlga* KO medaka showed the same phenotype, reinforcing the notion that *kitlga* is the causal gene of *fm* medaka.

Kit ligand, also known as stem cell factor (SCF), plays a key role in melanogenesis, gametogenesis, and hematogenesis in mammals (Copeland *et al*. 1990; Geissler *et al*. 1991). Homozygous mutation of the mouse Kit ligand gene {(*Kitl*)?} results in embryonic death due to severe macrocytic anemia, whereas heterozygous mutant animals are viable but manifest a wide spectrum of abnormalities including a variable extent of macrocytic anemia, a reduced number of mast cells, and reduced pigmentation including white spotting or a gray color of fur (Sarvella and Russell 1956; Broudy 1997). In mouse melanogenesis, melanoblasts are specified from neural crest cells, with *Mitf* and *Kit* being the earliest known markers for melanoblasts. After their differentiation, melanoblasts migrate dorsolaterally through the dermis between the somites and the developing epidermis from embryonic day 10.5 (Mort *et al*. 2015). Both *fm* mutant and our homozygous *kitlga* KO medaka are viable and manifest reduced numbers of melanophores and leucophores. This phenotype is thus similar to that of the heterozygous *Kitl* mutant mice, which have a reduced number of melanocytes.

Zebrafish has two *kitlg* genes, *kitlga* and *kitlgb*, with the former, but not the latter, playing a key role in the survival and migration of melanophores (Hultman *et al*. 2007). A zebrafish *kitlga* null mutant is viable and manifests a reduced number of melanophores, similar to our *kitlga* KO medaka. We found that medaka also harbors *kitlga* and *kitlgb* genes and that *kitlga* is the causal gene of the *fm* mutant, indicating that medaka *kitlga* is likely equivalent to zebrafish *kitlga*.

Tcb1 belongs to the Tc family of transposons and has been identified in nematodes and fruit flies. Transposons of the Tc family are ~1.6 kb in size, are associated with a TA repeat sequence, and contain a DDE motif in the open reading frame encoding the transposase (Harris *et al*. 1990; Hoekstra *et al*. 1999). Their inverted terminal repeats (ITRs) comprise 20 to 400 bp and contain CAGT at the 5’ end (Rosenzweig *et al*. 1983; Harris *et al*. 1988). The inserted sequence found in *kitlga* of *fm* medaka is similar to a Tcb1-like sequence found in other fish species. However, no ITR was associated with the Tcb1-like sequences detected in *fm* or WT medaka. Furthermore, this transposon-like sequence of medaka contains stop codons, indicating that the *kitlga* product in the *fm* mutant does not function like that in WT medaka. With the use of the dot plot program “dotmatcher” (http://www.bioinformatics.nl/cgi-bin/emboss/dotmatcher), we also did not find any other inverted repeats on either side of the inserted sequence in *fm* medaka, suggesting that the transposase is not active and that the insertion arose as the result of a “cut and paste” type mechanism.

c-Kit signaling activates the expression of Mitf via the Ras-Raf-Mek-Mapk and mechanistic target of rapamycin (mTOR) pathways in mammals, and Mitf promotes transcription of the gene for tyrosinase, which is the rate-limiting enzyme of melanin production (Rönnstrand 2004; Liang *et al*. 2013). Given that the number of melanophores is not reduced in the heterozygous *fm* mutant, partial loss of Kitlga production is likely not sufficient to result in down-regulation of the c-Kit signaling pathway. Melanophores of both *fm* medaka were found to be smaller than those of the WT at stage 36, whereas the size of leucophores or xanthophores in *fm* medaka did not differ from that in WT fish. The size of the remaining melanophores in *fm* embryos close to hatching was also previously found to be reduced (Kelsh *et al*. 2004). Furthermore, whereas melanophores were found to manifest an extended morphology beneath the epidermis in WT zebrafish larvae, they were rounded in the *spab5* mutant, which harbors a mutation in a *kit* ortholog (Parichy *et al*. 1999). These observations suggest that *kitlga* may contribute to the maturation or metabolism of melanophores as a result of activation of the expression of Mitfa by c-Kit signaling.

The genes *sox5*, *pax3a*, *pax7a*, *and mitf* play important roles in the development of chromatophores (Kimura *et al*. 2014; Nagao *et al*. 2014). We have now shown that the expression of these genes was suppressed in *kitlga* KO medaka, indicating that such expression is regulated by the Kitlga protein. Given that differentiation of all chromatophores from their multipotent progenitors is apparent in both *fm* and *kitlga* KO medaka, *kitlga* may influence the proliferation and migration of melanophores and leucophores after their differentiation. The development of iridophores and xanthophores is regulated by anaplastic lymphoma kinase (Alk) and leukocyte tyrosine kinase (Ltk) signaling and by colony-stimulating factor 1 (Csf1) signaling, respectively, in zebrafish (Patterson and Parichy 2013; Mo *et al*. 2017). These signaling pathways also may contribute to the proliferation and migration of the corresponding cell types in medaka (Figure S3).

Loss of function of c-Kit in humans gives rise to hypopigmentation-deafness disorders such as piebaldism and is also associated with certain tumor types such as thyroid carcinoma, melanoma, and breast cancer (Rönnstrand 2004; Dahl *et al*. 2015; Zazo *et al*. 2015; Tramm *et al*. 2016; Franceschi *et al*. 2017). Mice with white spotting also harbor heterozygous loss-of-function mutations in the c-Kit gene (Geissler *et al*. 1991). We did not detect tumors or organ abnormalities in either *fm* or *kitlga* KO medaka. Given that, as in the present study, changes in body coloration induced by drugs or genetic manipulation are readily detected in embryos or larvae of medaka within a matter of hours or days, our *kitlga* KO medaka may prove useful as a tool for screening of drugs for conditions related to loss of c-Kit signaling.

## ACKNOWLEDGMENTS

We thank S. Kuninaka for assistance with the generation of *kitlga* KO medaka as well as other laboratory members for assistance. This work was supported by the National Institute of Basic Biology (NIBB) Collaborative Research Program (c-17-312 and a-13-103).

---

**Figure S1** Molecular phylogeny of the *kitlg* gene in teleosts. The predicted amino acid sequences of *kitlg* genes of various species were downloaded from GenBank or Ensembl for construction of a phylogenic tree by the maximum likelihood method with 1000-bootstrap replication. Spotted gar was examined as an outlier. The numbers at each node indicate bootstrap probability. Gene names, species, GenBank accession numbers or Ensembl IDs, and amino acid sequences are provided in Table S1.

**Figure S2** Reduced expression of pigmentation-related genes in larvae of *fm* and *kitlga* KO medaka at 3 dph. Total RNA isolated from WT, *fm het*, *fm*, and *kitlga* KO larvae was subjected to RT and real-time PCR analysis of *pax7a* (A), *sox5* (B), *pax3a* (C), and *mitfa* (D) expression. Data were normalized by the amount of GAPDH mRNA and Data are means ± 95% confidence interval from three independent experiments. The *p* values were determined by one-way ANOVA followed by Tukey’s post hoc test.

**Figure S3** Model for chromatophore differentiation, proliferation, and migration from the neural crest. Melanophores and iridophores develop from a shared progenitor, as do xanthophores and leucophores. We propose that the proliferation and migration of leucophores and melanophores are regulated by c-Kit signaling, whereas the development of iridophores and xanthophores are thought to be regulated by Alk-Ltk signaling and Csf1 signaling, respectively.

**Figure s1.**
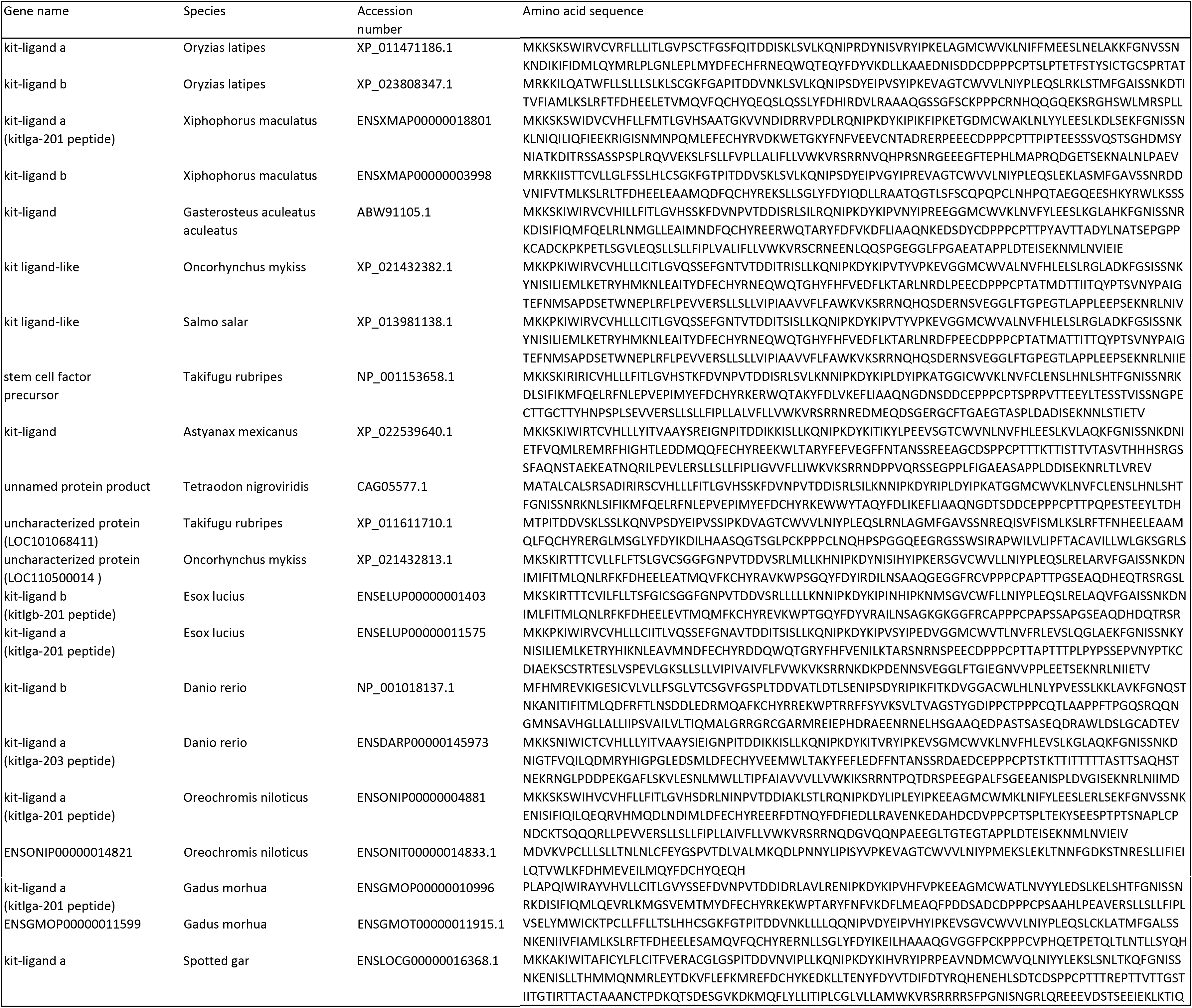
The genome list in the phylogenetic tree

